# Melatonin reduces GSK3β expression and tau phosphorylation *via* Nrf2 nuclear translocation

**DOI:** 10.1101/861229

**Authors:** Rashmi Das, Abhishek Ankur Balmik, Subashchandrabose Chinnathambi

**Author notes:** The authors wish it to be known that, in their opinion, the first two authors should be regarded as joint First Authors. To whom correspondence should be addressed**: Prof. Subashchandrabose Chinnathambi**, Neurobiology group, Division of Biochemical Sciences, CSIR-National Chemical Laboratory (CSIR-NCL), Dr. Homi Bhabha Road, 411008 Pune, India. **Abbreviation.** AD, Alzheimer’s Disease; APP, Amyloid precursor protein; Aβ, Amyloid-β; NFTs, neurofibrillary tangles; IL, Interleukin; CNS, Central nervous system; ROS, Reactive oxygen species; MAP3K, MAP kinase kinase kinase; Nrf2, Nuclear factor erythroid-2 related factor 2; pGSK3β, Phosphorylated glycogen synthase kinase 3β; TNF α, Tumor necrosis factor α; TGF β, Tumor growth factor β.

## Abstract

Alzheimer’s disease is a neuropathological condition with abnormal formation of extracellular Amyloid-β plaques and intracellular neurofibrillary tangles (NFTs) of Microtubule-associated protein Tau (Tau) in brain. In pathological condition, MAP-Tau can undergo various post-translational modifications such as hyperphosphorylation by the activity of cellular kinases which eventually leads to protein aggregation in neurons. Melatonin is a hormone which mainly secreted from pineal gland, functions to modulate cellular kinases. In our study, we elucidated that Melatonin has inhibited the Tau aggregates mediated cytotoxicity and membrane leakage by MTT and LDH assay respectively in neuro2A cells. Melatonin has found to reduce the GSK3β mRNA expression and protein level by western blot and immunofluorescence assay. Melatonin has also decreased phospho-Tau level (pThr181 and pThr212-pSer214) in neuron cell line upon OA induction as seen by microscopic analysis.. Melatonin treatment has associated with ROS quenching by DCFDA assay, reduced caspase 3 activity in neuronal cells. Further, Melatonin has increased Nrf2 level and nuclear translocation as oxidative stress response in Tauopathy. Together, these findings clearly signifies that Melatonin remediate the Tau-induced neuronal cytotoxicity and reduce Tau hyperphosphorylation *via* downregulating GSK3β expression. Melatonin can combat oxidative damage by Nrf2 activation and nuclear translocation in AD condition.

## INTRODUCTION

Alzheimer’s Disease (AD) is a chronic neurodegenerative disease which is associated with extracellular and intracellular deposition of β-amyloid plaques and NFTs of MAP-Tau in CNS. Tau is a natively unfolded cytosolic protein which functions in microtubule stabilization and axonal transport (Johnson and Hartigan, 1999). Tau is found to be post-translationally modified of which phosphorylation is the most common in AD. Tau becomes phosphorylated by various cellular kinases (GSK3β, CDK5-P25, PKA). The pThr231 (PHF6), pSer202-pThr205 (AT8), pSer212-pThr214 (AT100), pThr181 (AT270) phosphorylation of Tau are mainly found in Tauopathy (Mandelkow and Mandelkow, 2012). Phosphorylated Tau is also crucial in developmental regulation as evident from fetal Tau which is more phosphorylated than adult Tau in brain. Age-related dephosphorylation of Tau by phosphatases are witnessed in neurodegenerative diseases. For example, binding of PP2A and PP2B causes dephosphorylation of Serine 396 residue, alteringthe microtubule-binding function of Tau (Matsuo et al., 1994).

Melatonin is N-acetyl 5-methoxy tryptamine, a small hormone mainly produced from pineal gland which regulates the sleep-awake cycle (Claustrat et al., 2005). It functions as free radical scavenger, anti-apoptotic, anti-cancer, neuroprotectant and immune modulator. (Balmik and Chinnathambi, 2018). Melatonin can also induce the expression of nuclear factor erythroid-2 related factor (Nrf2) to reduce the oxidative burden in rat urinary bladder cells (Tripathi and Jena, 2010). Melatonin treatment has been shown to improve cognitive function (Lin et al., 2013), lessen aggregates burden and sleeplessness in neurodegenerative diseases (Quinn et al., 2005). Melatonin stalls the aggregation of β-amyloid peptide and inhibits the production of pro-inflammatory cytokines (IL 1, IL 6, TNF α) in β-amyloid rat model (Rosales-Corral et al., 2003). Melatonin interferes with the inflammatory process by interacting with transcription factor NFκB and iNOS activity was found to be decreased in J774 and RAW 264.7 murine macrophages (Gilad et al., 1998). Additionally, Melatonin involved in minimizing the production of endothelial adhering molecules (ICAM), vesicular adhering molecule (VCAM-I), endothelial selectin (E-selectin), releated to reduced inflammatory burst (Motilva et al., 2011). Melatonin ameliorates the amyloid-β overload via PI3-Akt signaling and ultimately reduces the Tau hyperphosphorylation in mouse hippocampus (Ali and Kim, 2005).

In this study, we investigated the protective role of Melatonin in Tau mediated neurodegeneration and its potentiality to reduce the oxidative damage and apoptotic loss in Tauopathy. We identified the efficacy of Melatonin in the reduction of Tau phosphorylation, GSK3β expression level and Nrf2 translocation during oxidative stress condition during Tau overloaded inflammatory burst.

## MATERIALS AND METHODS

### Cell Cultures and reagents

Luria-Bertani broth (Himedia); Ampicillin, NaCl, Phenylmethylsulfonylfluoride (PMSF), MgCl_2_, APS, DMSO, Methanol, Ethanol, Chloroform, Isopropanol were purchased from MP Biomedicals; IPTG and Dithiothreitol (DTT) from Calbiochem; MES, BES, SDS, MTT reagent, Melatonin, 2’-7’-Dichlorodihydrofluorescein diacetate (DCFDA), EnzChek^™^ Caspase-3 Assay Kit (Z-DEVD-AMC substrate) (E13184), Okadaic acid, TritonX-100, alpha Actin Loading Control Monoclonal Antibody (A2066) from Sigma; EGTA, Protease inhibitor cocktail (PIC), Tris base, 40% Acrylamide, TEMED, Dulbecco modified eagle’s media (DMEM), Fetal bovine Serum (FBS), Horse serum, Phosphate buffer saline (PBS), trypsin-EDTA, Penicillin-streptomycin, Pierce^™^ LDH Cytotoxicity Assay Kit (88953), RIPA buffer, mouse Beta Tubulin (BT7R) (MA516308),, mouse Tau antibody T46 (13-6400), AT100 (MN1060), Phospho-Tau (Thr181) Antibody (5H9L11) Rabbit anti GSK3β monoclonal antibody (PA5-68817), Rabbit anti p-GSK3β (Ser9) monoclonal antibody (MA515109), Rabbit anti Nrf2 polyclonal antibody (PA5-68817), Goat anti-Rabbit IgG (H+L) Cross-Adsorbed Secondary Antibody HRP (A16110), anti-mouse secondary antibody conjugated with Alexa flour-488 (A-11001), Goat anti-Rabbit IgG (H+L) Cross-Adsorbed Secondary Antibody with Alexa Fluor 555 (A-21428), DAPI, Trizol reagent, First strand cDNA synthesis kit (K1612), DNA oligonucleotide primers from IDT, Maxima SYBR Green/Fluorescein qPCR Master Mix (2X) (K0241) from Invitrogen. The coverslip (0.17 mm) was from Blue star for immunofluorescence study.

### Preparation of Tau aggregates and Transmission Electron Microscopy

The full-length human Tau (4R2N) was expressed in *E.coli* BL21* in presence of 100 μg/ml of ampicillin. The protein purification was done as described previously (Gorantla et al., 2017). Tau aggregates were prepared by inducing Tau aggregation with Heparin (17.5 kDa) in 20 mM BES buffer, pH 7.4 (Heparin/Tau ratio-1:4) and incubated for 7 days at 37°C (Barghorn et al., 2005). Formation of mature fibrils were checked by pelleting assay at 60000 rpm for 60 minutes, followed by SDS-PAGE and Transmission electron microscopy (TEM) with 120 kV electron beam.

### Cell viability assay

Neuro2A cell line was cultured in DMEM with 10% FBS and 100 μg/mL of penicillin-streptomycin. Ten thousand cells/well were seeded in 96-well plates. Melatonin was added at different concentrations, with 5% DMSO as a control at final volume of 100 μL. Tau aggregates (10 μM) were added separately and along with various concentrations of Melatonin (from 0.1 to 100 μM) and incubated for 24 hours at 37°C. MTT (Thiazolyl blue tetrazolium bromide) reagent was added at final concentration of 0.5 mg/mL in each well and incubated for 3 hours at 37°C. The formazan end product was solubilized by adding DMSO and the absorption was measured at 570 nm in spectrophotometer (Infinite® 200 M PRO, Tecan).

### Lactate Dehydrogenase (LDH) assay

Aggregates induce cellular toxicity and alter membrane permeability which leads to the leakage of cytosolic enzyme LDH(Flach et al., 2012). Neuro2A cells were treated with 10 μM of aggregated Tau and different concentrations (from 1 to 50 μM) of Melatonin together for 24 hours. The cell-free media were collected from each wells and LDH leakage was measured by the formation of formazan compound which is directly proportional to the cytotoxicity as per manufacturer’s protocol (Pierce^™^ LDH Cytotoxicity Assay Kit).

### Detection of intracellular ROS production by DCFDA assay

In order to check intracellular ROS production by Tau aggregates, neuro2A cells were treated individually and in combination with 1 and 10 μM concentrations of Melatonin and 10 μM of Tau aggregates for 1 hour along with 5% DMSO as positive control. Then, the cells were washed thrice with PBS and incubated with 25 μM DCFDA for 30 minutes. Intracellular ROS cleaves DCFDA and the amount of DCF fluorescence is directly proportional to ROS production which was detected by Flow cytometry (BD Accuri C6). The fluorescence positive cell population was selected by live-dead gating (>80% from the whole population) and then acquire 50,000 cells through FITC channel (Excitation/Emission: 488/530 nm). The relative fluorescence of the test groups were calculated by subtracting the values of autofluorescence of untreated cell control.

### Caspase assay

Caspase-3 activation can be quantified by fluorometric assay in which a signal peptide is linked with a fluorophore molecule (DEVD-Rhodamine110). Okadaic acid (OA) is an inducer of most cellular protein phosphorylation and apoptosis, leading to neuronal death(Lavrik et al., 2005; Suuronen et al., 2000). Neuro2A cells were treated with Tau aggregates (10 μM) and OA (25nM) with Melatonin (50 μM), and incubatedfor 6 hours. Then, the cells were harvested, lysed and mixed with fluorophore conjugated signal peptide substrate and incubated for 30 minutes. Caspase 3 cleaves the signal peptides and the amount of released fluorophores is measured by Excitation/Emission at 485/535 nm which is proportional to the apoptotic activity.

### Western Blot

To study the over-activation of GSK3β, we treated the neuro2A cells with okadaic acid (25 nM) and Melatonin (50 μM) separately and together for 24 hours. The cells were washed with PBS and lysed with RIPA buffer and cell lysate were subjected to western blot with anti GSK3β monoclonal antibody (1:2500), anti p-GSK3β monoclonal antibody (1:2000) and anti Nrf2 polyclonal antibody (1:1000) with β-tubulin (1:5000), α-Actin (1:2500) as internal control. Then, the bands intensity were quantified by using BIORAD Quality one 4.6.6 software. The band-density of the treated group were compared with its corresponding untreated control group and normalized with specific house-keeping gene control (β-tubulin or α-Actin) (n=2). Then, the relative fold changes were plotted with respective proteins and treated group.

### Immunofluorescence Assay

The level of Tau phosphorylation upon okadaic acid (25 nM) treatment and the role of Melatonin (50 μM) on cellular kinase-p-GSK3β (Ser9), were checked by Immunofluorescence study along with oxidative stress response transcription factor Nrf2. Neuro2A cells were treated with these compounds together and separately for 24 hours. Then, the cells were washed with PBS thrice and fixed with chilled absolute methanol for 15 minutes and permeabilized with 0.2% TritonX-100. The cells were stained with Phospho-Tau (AT100 1:100), Phospho-Tau (pTau (Thr181) 1:100), p-GSK3β (Ser^9^) (1:100), GSK3β (1:100), Nrf2 (1:100) and T46 (1:250) antibody for overnight in 2% horse serum. Then, Alexa flour-secondary antibody was allowed to bind p-Tau, p-GSK3β, T46, total GSK3β and Nrf2 along with total Tau for 1 hour along with nuclear stain-DAPI (300 nM). The microscopic images were taken in Zeiss Axio observer with Apotome2 fluorescence microscope at 63X oil immersion. The quantification were done using Zen2 software and the mean fluorescence intensity was plotted for different test groups.

### Expression profile study by Quantitative real-time PCR

Neuro2A cells were treated with Melatonin (50 μM) and okadaic acid (25 nM) for 6 hours. Then the cells were washed PBS and total RNA were isolated by the conventional TRIZOL reagent, chloroform and isopropanol extraction procedure. The RNA pellets were washed with 80% ethanol, dissolved in DEPC treated water, and proceeded for cDNA synthesis (First-strand cDNA synthesis kit) using oligo-dT primer. The expression of protein kinases (GSK3β, MAP3K) and NFkB component (P65) were checked by qRT-PCR (Table 1). The fold change was calculated by ΔΔCT method with respect to house-keeping GAPDH control.

**Table.**
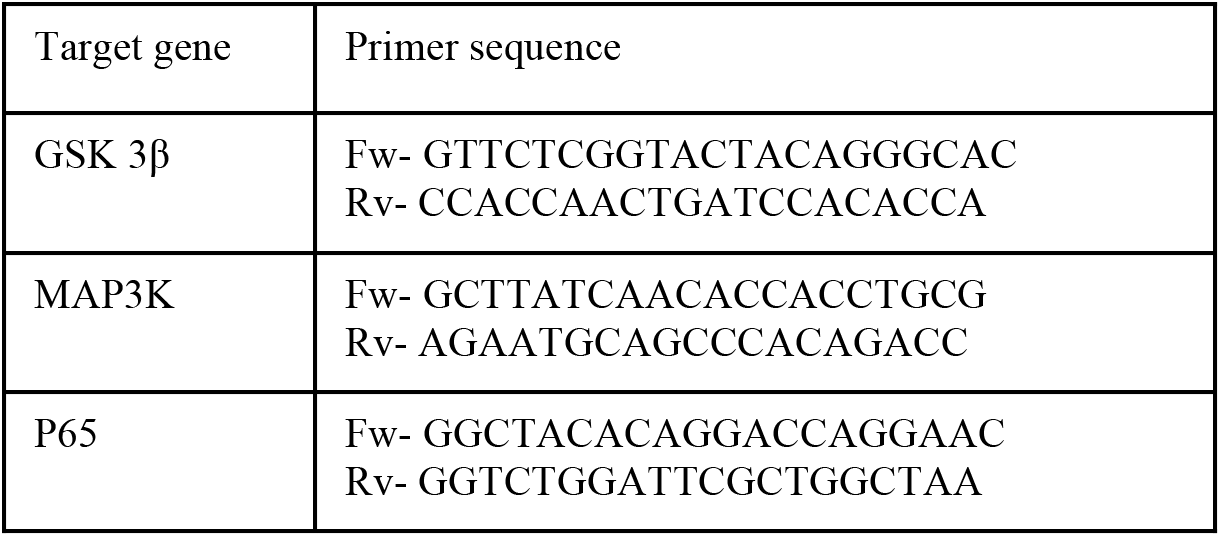
Primer list used for qRT-PCR expression profile study.

### Statistical Analysis

Each experiment was repeated for three times and the measurements for every experiments were taken in triplicate. Statistical analysis were performed by using two-tailed student’s t-Test in SigmaPlot 10 software. Okadaic acid and Melatonin treated groups were compared with untreated cell control while ‘okadaic acid and Melatonin together group was compared with only okadaic acid-treated group. p-values were calculated and represented as p<0.05 as *, p<0.001 as ** and p<0.0001 as ***.

## RESULTS

### Melatonin reduces Tau-mediated cytotoxicity

Extracellular deposition of amyloid-β and intracellular NFTs of Tau induce neuronal toxicity, lead to cognitive impairment due to synaptic loss in AD. Melatonin is known as neuroprotectant, improving mitochondrial functions, antioxidant, immune modulator (Galano et al., 2014; Reiter et al., 2016) and restore cognitive function (Cardinali et al., 2012). In our study, Melatonin has reduced the Tau mediated toxicity in a concentration-dependent manner. When neuro2A cells were treated with only Melatonin, it showed 50% viability at 1 mM concentration by MTT assay (Fig. 1A) and phase contrast study (Suppl. Fig. 1A). Upon treatment of 10 μM of Tau aggregates, neuro2A cells have shown 60% viability but, Melatonin at 20 μM concentration has rescued the viability up to 80% in Tau-stressed condition (Fig. 1B). The cytotoxicity was completely reversed at 200 μM concentration of Melatonin (Suppl. Fig. 1A).

**Figure 1.**
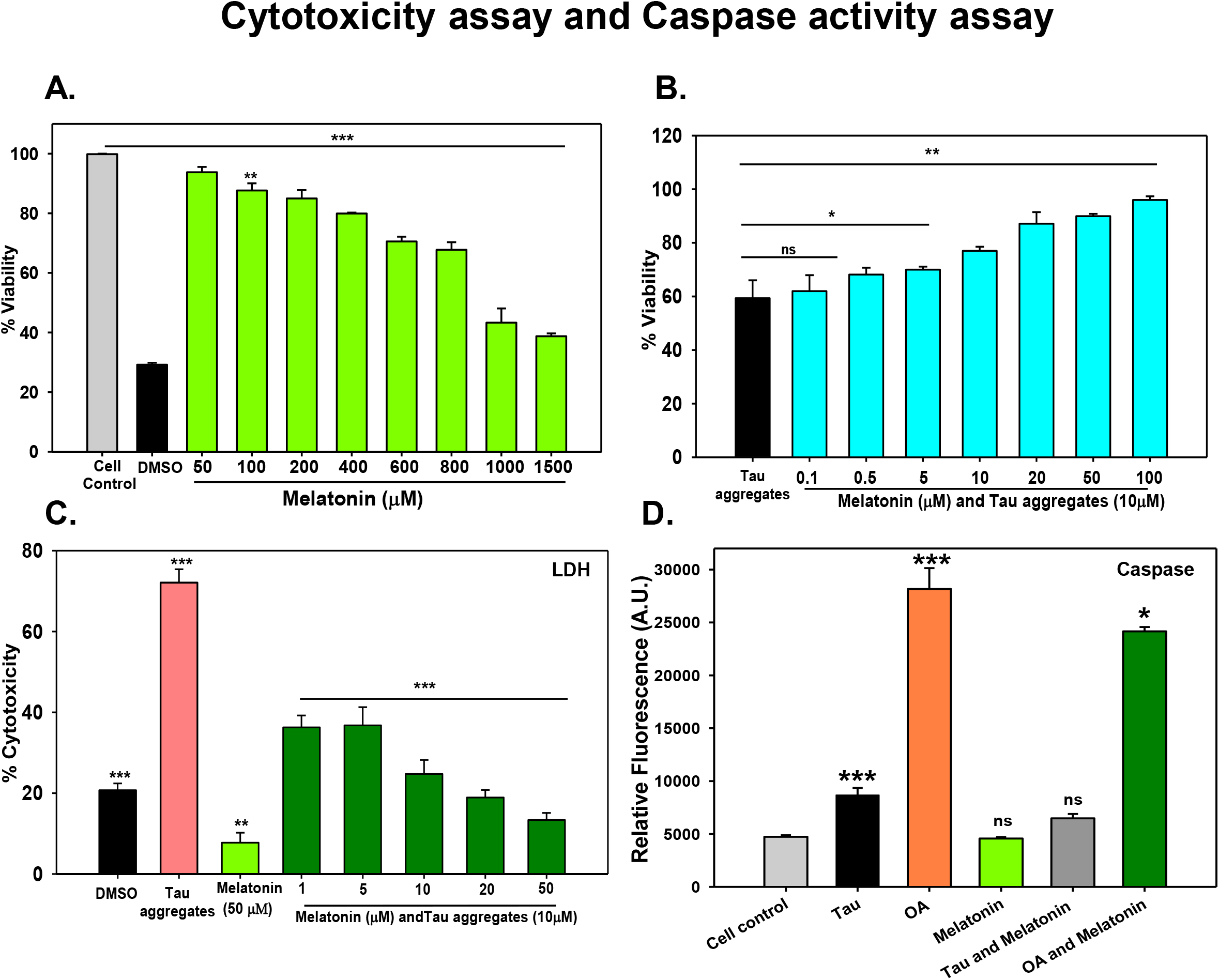
Cytotoxicity and caspase activity assay. (A) Melatonin showed cytotoxicity at only higher concentrations (>1 mM) on neuro2A cells as found by MTT formazon production. (B) Pre-formed Tau aggregates at 10 μM showed 60% viability on neuro2A cells, but when the cells were treated along with Melatonin, the cell viability increased in a concentration dependent manner. (C) Cytotoxicity study was confirmed by LDH assay. Tau aggregates (10 μM) induced 70% cytotoxicity whereas Melatonin treatment reduced the Tau aggregates mediated cytotoxicity in a concentration dependant way. (D) OA induced high caspase activity as compared to Tau aggregates treatment. Melatonin treatment reduced the OA induced caspase activity effectively but Tau mediated caspase activity remain unaltered. Values were given as mean + SEM. * corresponds to test groups compared with untreated control (* p < 0.05; ** p <0.01, *** p<0.001).

The leakage of cytoplasmic enzyme LDH is a determinant of cytotoxicity and membrane damage. Tau aggregates treatment on neuro2A cells has resulted in 68% cytotoxicity while the 10 μM concentration of Melatonin has reduced 50% Tau mediated cytotoxicity (Fig. 1C). Pelleting assay, SDS-PAGE analysis have showed the formation of mature Tau fibrils with heterogeneous mixture of aggregates of 150-250 kDa and fibrillar aggregates as seen by TEM analysis (Suppl. Fig. 1B, C).

### Anti-apoptotic role of Melatonin

Caspases-3 is the central effector for the aspartate-guided cleavage of the target protein in both intrinsic (mitochondrial) and extrinsic (DEAD ligand) apoptotic pathway. Previous report suggested that caspase-3 can cleave APP which leads to further aggregation (Gervais et al., 1999). Melatonin has reported to prevent the intrinsic pathway of apoptosis in neurodegenerative diseases (Wang, 2009). Here, Tau aggregates and OA have induced two fold and five fold caspase 3 activity respectively on neuro2A cells as compared to untreated control. Melatonin has reduced the OA-mediated caspase activation at 50 μM concentration while Tau aggregates mediated caspase activation remain unaltered by Melatonin (Fig. 1D).

### Melatonin has reduced GSK3β level but not p-GSK3β (Ser^9^)

Modified Tau dissociates from microtubules and become aggregated in cytosol during AD (Avila, 2006). GSK3β, a MAPK family kinase, with altered activation state at Ser9 phosphorylation, mediates hyperphosphorylation of Tau in AD (Sun et al., 2015). To test the role of Melatonin on GSK3β-mediated Tau-phosphorylation (Fig. 2A), neuro2A cells were treated with OA for induction of phosphorylation, followed by western blot and immunofluorescence with phospho-GSK3β (Ser^9^) and total GSK3β antibody. OA has upregulared the mRNA expression of GSK3β whereas, Melatonin significantly downregulated the GSK3β expression in OA-stressed neuronal cells (Fig. 2B). The global phosphorylation has induced by OA as seen in higher p-GSK3β (Ser^9^) level, compared to total GSK3β level (Fig. 2C, E). Melatonin treatment on neuro2A cells has reduced the total GSK3β protein level but the phosphorylation at Ser^9^ of GSK3β as an altered activated state remained invariable (Fig. 2D). Immunofluorescence study depicted the upregulation of total GSK3β and phospho-GSK3β level in OA exposure (Fig. 2F) but only total GSK3β level was reduced in melatonin-treated and OA-stressed neurons (Fig. 2G, H).

**Figure 2.**
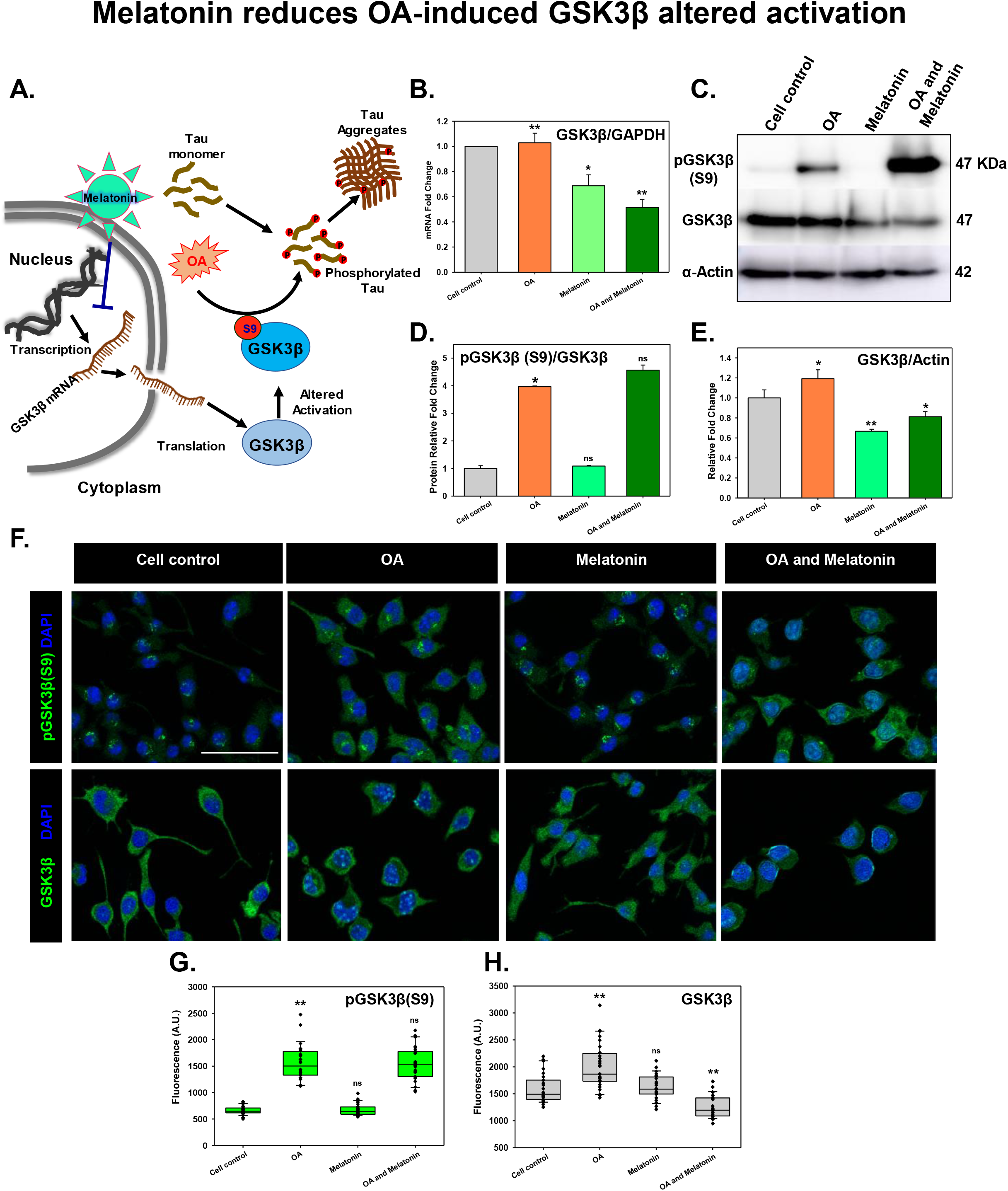
Melatonin reduces OA-induced GSK3β altered activation. (A) During AD, Tau become hyperphosphorylated and subsequently aggregation in neurons. OA has been proven as a model to induce Tau phosphorylation *via* altered activation of GSK3β (phopsho Ser^9^). Melatonin has been proposed to block the GSK3β-mRNA expression and total protein level. (B) Melatonin has reduced the total-GSK3β mRNA expression level on neuro2A cells upon OA treatment. (C, D) OA has induced the Ser^9^ phosphorylation of GSK3β as altered activation state whereas the total GSK3β has been downregulated. (E) Western blot densitometric quantification showed the decreased level of total GSK3β. (F) Immunofluorescence study depicted the altered level of only total GSK3β protein but GSK3β (phopsho Ser9) remained invariant. (G, H) The relative fluorescence level were plotted for Melatonin and OA treated group alone and together. * corresponds to test groups compared with untreated control (* p < 0.05; ** p <0.01, *** p<0.001). scale bar: 50 μm.

### Melatonin decreases pTau level

Tau has undergone hyperphosphorylation by the overactivation of cellular kinases in AD and thereby looses its affinity towards microtubule and remains freely diffusible in cytosol. pTau forms oligomeric seed species by lowering its intermolecular interaction and aggregated as insoluble inclusions. Altered activated-GSK3β(Ser^9^) induces the phosphorylation of Tau at different Serine/Threonine residues (Yuan et al., 2004) and changes its subcellular localization from axons to cell bodies, even at nucleus in AD (Lu et al., 2013). Melatonin depleted the pTau (Thr181) level in neuro2A cells which was found to be localized into the nucleus (Fig. 3A). But the distribution of nuclear-pTau (Thr181) became distorted upon OA exposure and observed to be concentrated in nuclear periphery (Fig. 3B, C). Phosphorylation at AT100 epitope of Tau was observed to localize around the cytoplasmic membrane, which was more amplified in OA-treated cells and less in Melatonin exposure (Fig. 3D).

**Figure 3.**
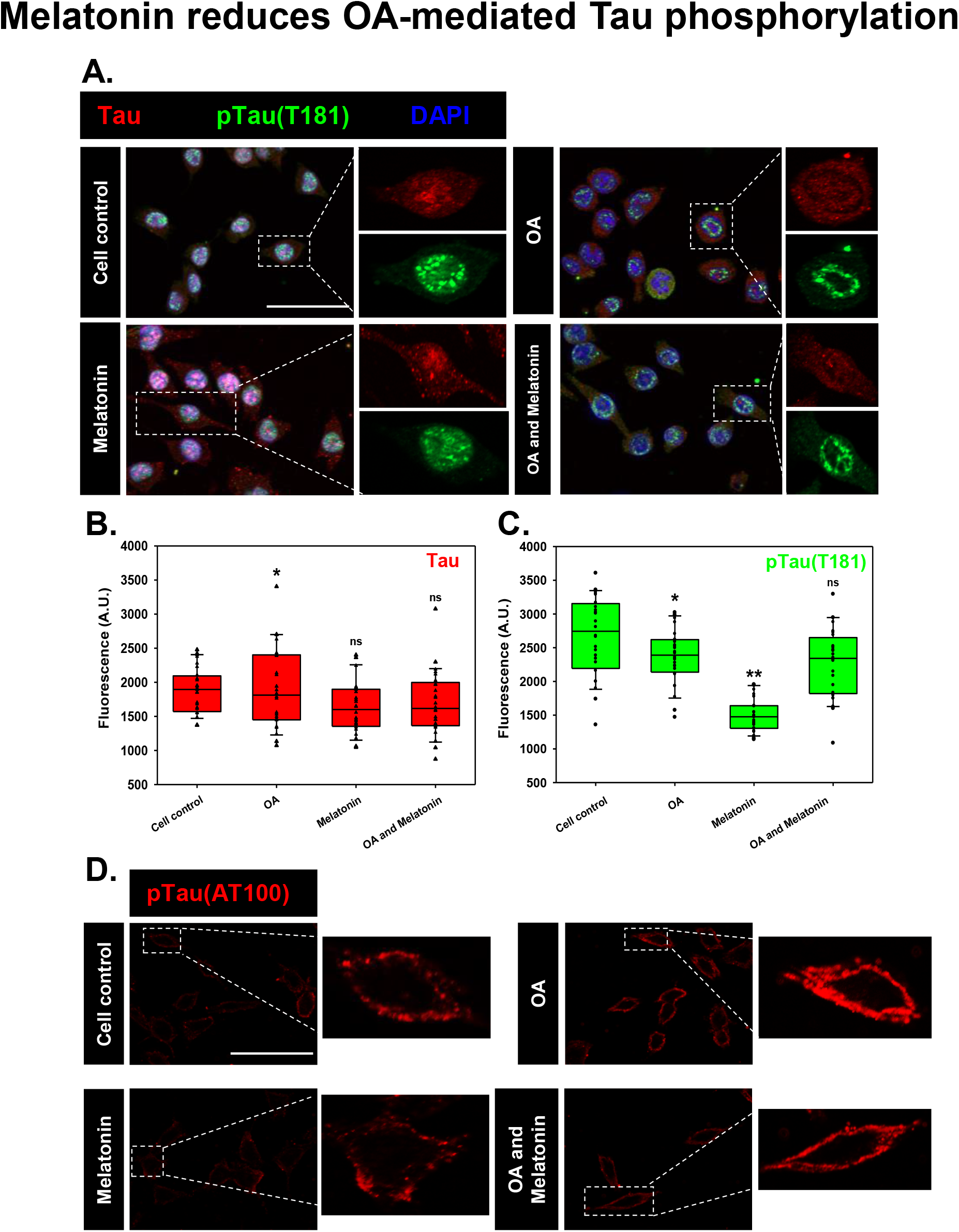
Melatonin reduces OA-mediated Tau phosphorylation. (A) Phopho-Tau (Thr181) is one of the epitopes, phosphorylated by GSK3β in AD condition. Phopho-Tau (Thr181) has found to be localized into the nucleus while OA exposure has concentrated the particular phospho-Tau epitope only in the nuclear periphery. (B, C) Mean fluorescence quantification has showed the decrease of Phopho-Tau (Thr181) level by Meltonin treatment. (D) Phopho-Tau (AT100) has been shown to localize in the membrane periphery, which was induced by OA. Melatonin rescued the AT100-phosphoTau level in OA-stressed neuro2A cells. scale bar: 50 μm.

### Melatonin quenches intracellular ROS production and mediates Nrf2 translocation into nucleus

Melatonin is known as antioxidant molecule, which not only scavenges reactive oxygen but also modulates the activity of antioxidant enzymes like glutathione peroxidase, glutathione reductase, superoxide dismutase *etc*. (Zhang and Zhang, 2014). Melatonin plays a crucial role in mitochondrial homeostasis and stabilizes electron transport chain, ATP production and mitochondrial transportation (David et al., 2005; López et al., 2009). Tau treatment has increased the intracellular ROS level to 700 A.U. which was similar to DMSO of ~500 A.U., as observed by DCFDA fluorescence. Melatonin at 10 μM concentration has reduced almost 50% of ROS level on Tau-overloaded neuro2A cells (Fig. 4B). As evidenced, Melatonin has ROS scavenging property and has the potentiality in rescuing the neurons from Tau stress mediated oxidative damage.

**Figure 4.**
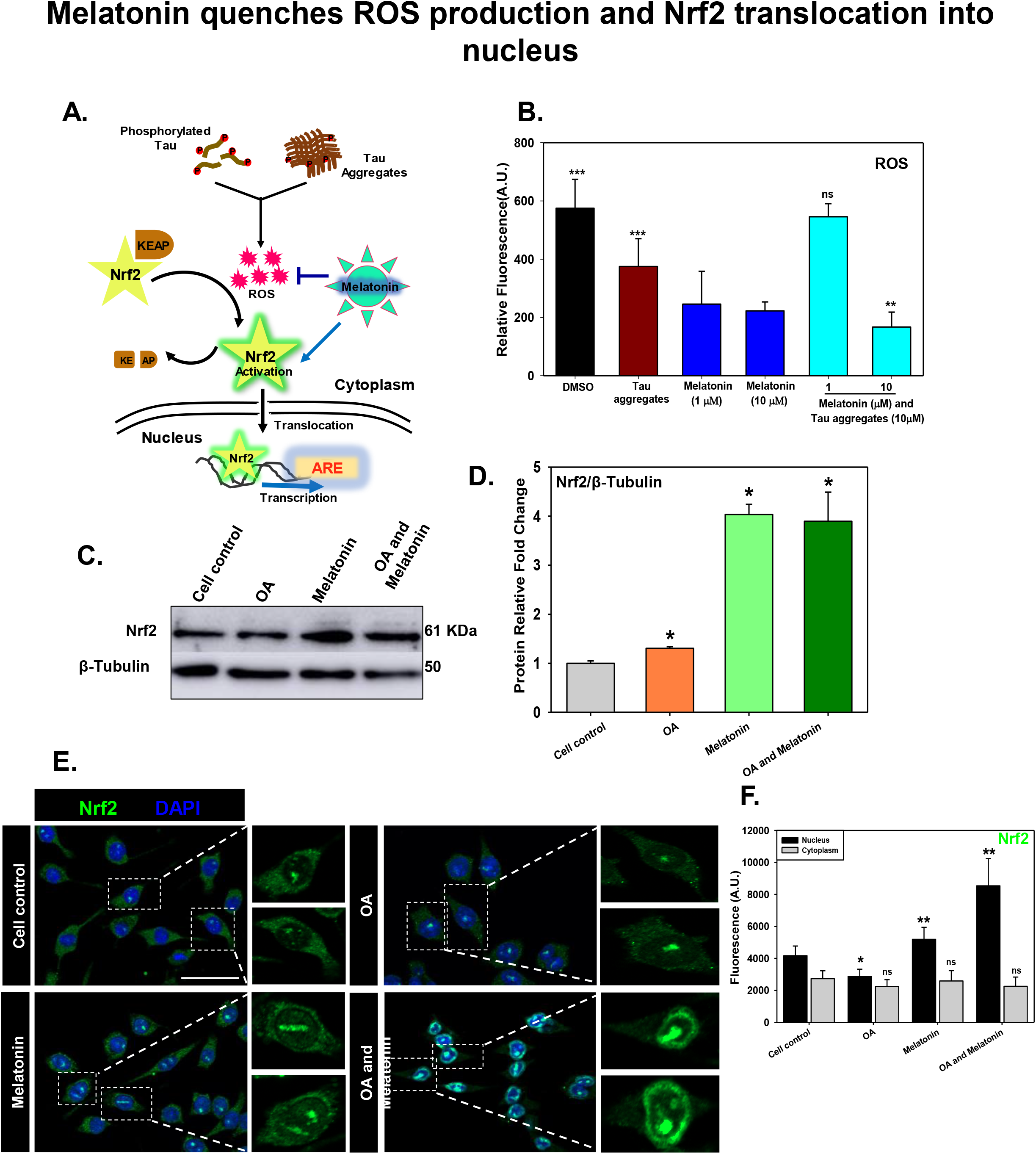
Melatonin quenches ROS production and Nrf2 translocation into nucleus. (A) Intracellular phosphorylated Tau and Tau aggregates have the potentiality to induce the ROS production. Nrf2-KEAP1 complex presents in cytosol as inactive form, but oxidative stress induces the dissociation of KEAP from Nrf2. Free Nrf2 become activated and translocated into the nucleus for transcription of anti-oxidant responsive element (ARE) as global production of anti-oxidant enzymes and subsequent impeled proteostasis. (B)Tau aggregates treatment induced intracellular ROS level while Melatonin at 10 μM concentration has reduced the intracellular ROS level (DCFDA fluorescence) significantly as detected by FACS. (C, D) Melatonin rescued the neurons from oxidative damage by increasing the protein level of Nrf2 extensively as compared with β-tubulin as loading control in OA-stressed neuro2A cells. (E, F) Melatonin treatment has associated with induced Nrf2 level into nucleus while the complete nuclear translocation was evidenced alonh with OA-streesed condition by immunostaining. scale bar: 50 μm.

**Figure 5.**
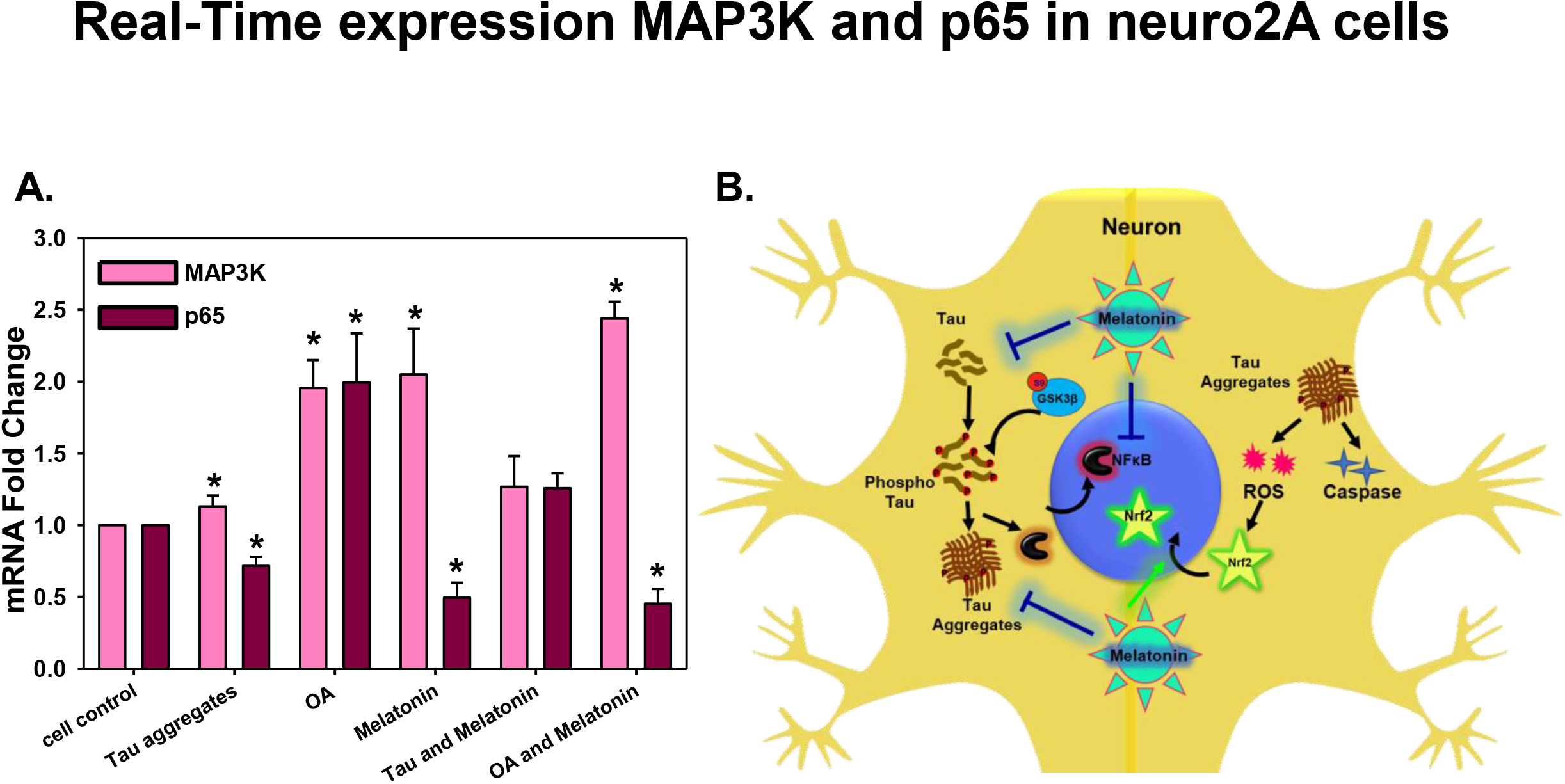
Multifactorial function of Melatonin in Tau-mediated neurodegeneration. (A) mRNA expression level of p65 (NFκB) increased upon Tau and OA treatment whereas; Melatonin reduced the P65 level compared to untreated group. MAP3K expression level was increased by 2.5 times upon Melatonin treatment, related to neuronal survival. Values were given as mean + SEM. * corresponds to test groups compared with untreated control (* p < 0.05; ** p <0.01, *** p<0.001). (B) Activation of cellular kinase leads to Tau phosphorylation and aggregates accumulation, which in turn activates ROS production and Caspase-apoptotic signal. Aggregated Tau can be released through axons, which are eventually uptaken by microglia and returned to ‘Activation’ state. Activated microglia enlarges its extension and produce excess pro-inflammatory cytokines. But Melatonin treatment reduce the activated state of microglia and allows to secrete more anti-inflammatory cytokines, which can manage the burden of aggregates by increased phagocytic activity. Melatonin reduces NFκB activation in inflammation, but MAP3K activation and the transition towards M2 phenotype of microglia leads to axonal repair and survival.

Nrf2 is an essential transcription factor, which induce the expression of anti-oxidant mediators. Under physiological condition Nrf2 is a short lived protein, but upon activation Nrf2 become stabilized and translocated to the nucleus and subsequently activates the transcription of anti-oxidant enzymes and anti-inflammatory genes (Fig. 4A). In western blot study, Melatonin has induced Nrf2 expression even in presence of OA stress which clearly signifies that Melatonin is directing through Nrf2-mediated anti-oxidant pathway in neurons (Fig. 4C, D). In immunofluorescence study, Nrf2 level was found to be increased and localized into nucleus by Melatonin treatment. In OA-stressed neurons, Nrf2 was completely translocated into nucleus upon Melatonin treatment, which clearly emphasizes on oxidative stress response and corresponding rescuing effect by Melatonin (Fig. 4E, F).

## DISCUSSION

AD is a multi-factorial disease which includes the aggregation of β-amyloid protein as plaques in brain. An altered signaling cascade leads to abnormal modifications of intracellular protein-Tau which leads to NFTs formation (Mandelkow and Mandelkow, 2012). Excessive aggregates affect the neuro-trafficking, neurotransmitter release, leading to axonal blockage and attenuating neuronal plasticity (Guo and Lee, 2014). Exposure of the protein aggregates activate microglia with increased production of ROS/NO (Bedard and Krause, 2007), inflammatory cytokines, chemokines and complements (Schetters et al., 2018) which, ultimately results in the loss of neuronal synapses by phagocytosis (Mirbaha et al., 2018). Neurological disorders are often associated with impaired circadian rhythm due to the shortage of pineal gland produced Melatonin level (Zhou et al., 2003). Since last decades, Melatonin has been identified as a mighty lead as natural anti-oxidant, anti-cancer, anti-inflammatory molecule, enzyme regulator, metabolic-energy sustainer, epigenetic coordinator and aggregation inhibitor (Balmik and Chinnathambi, 2018). Here, we found that Melatonin has reduced the Tau aggregates-mediated neuro-toxicity at low concentration and remain safe even at higher concentration. Membrane leakage by the Tau fibrils and subsequent spreading from affected neurons are very much prominent events in Tauopathy (Guo and Lee, 2014). In our study, we evidenced the initiation of membrane leakage on neurons upon Tau fibrils exposure. But the event was restrained by Melatonin, which signifies its role in remodeling of membrane integrity in AD. We perceived that Melatonin can prevent the OA-induced apoptosis in neurons but when Tau aggregates were populated, Melatonin can not reduce the caspase activation. Hyperphosphorylated Tau is one of the pathological consequences of AD, which is mediated by the over-activation of cellular kinases(Sun et al., 2015). Melatonin regulates kinase activity by interfering with intermediate signaling cascade Akt/PI3K (Ali and Kim, 2005) or receptor-mediated mechanism (Wang et al., 2004).

Phosphorylated Tau were co-localized with Aβ on synaptosomal membrane compartments (Fein et al., 2008). Our study showed that Melatonin can reduce the level of nuclear as well as membrane-associated phospho-Tau (pTau181 and AT100 epitope) by downregulating the GSK3β transcription and hence, protein level. But, the altered activation of GSK3β at pSer^9^ which acts as a key kinase in Tau-phosphorylation (Yuan et al., 2004), can not be minimized by Melatonin.

Being an anti-oxidant transcriptional activator, Nrf2 functions to expand the level of different enzymes like-catalase, SOD, peroxidase, detoxifying enzyme in xenobiotic metabolism to oxidative stress (Murphy and Park, 2017). Nrf2 also play a crucial role in microglial phenotypic determination. Depletion of Nrf2 has induced the inflammatory burst, NFκB activation and reduced phagocytic activity in primary microglia upon α-synuclein exposure. Nrf2^−/−^ mice have shown an impaired ubiquitin-proteasomal system in α-synuclein overload as compared to WT (Lastres-Becker et al., 2012). Hence, the activation of global antioxidant transcription factor-Nrf2 along with neurtropic factor can be a potential therapeutic strategy to cambat AD (Murphy and Park, 2017). We found that Melatonin has quenched ROS production and mediates anti-oxidation *via* activation of Nrf2 and subsequent nuclear translocation.

All Together, Melatonin plays combinatorial role in Tauopathy, by reducing Tau phosphorylation, dissoluting aggregates-mediated toxicity, reducing membrane leakage in neurons. It also mediates the anti-oxidant and anti-apoptotic function by quenching free radicals, deactivating Caspase-3. Melatonin has also rescued OA-induced Tau phosphorylation by downregulating GSK3β expression and Nrf2 activation-translocation. So, the Melatonin conjugated with conventional drug can be proposed for multifunctional approaches in the treatment of AD.

## FOOTNOTES

## Acknowledgements

This project is supported in part by grants from the Department of Biotechnology, Neuroscience Task Force (Medical Biotechnology-Human Development & Disease Biology DBT-HDDB))-BT/PR/19562/MED/122/13/2016 and in-house CSIR-National Chemical Laboratory grant MLP029526. Tau constructs were kindly gifted by Prof. Roland Brandt from University of Osnabruck, Germany. The authors acknowledge Dr. H V Thulasiram for q-real time PCR facility. Special thanks to Mr. Ashish Kumar for helping in expression profile study. Special thanks to Mr. Tushar Dubey for proof-reading the article.

Rashmi Das acknowledges the fellowship from University Grant Commision (UGC) India. Abhishek Ankur Balmik acknowledges Shyama Prasad Mukherjee fellowship (SPMF) from Council of Scientific Industrial Research (CSIR), India.

